# Use of a Neural Circuit Probe to Validate in silico Predictions of Inhibitory Connections

**DOI:** 10.1101/204594

**Authors:** Honglei Liu, Daniel Bridges, Connor Randall, Sara A. Solla, Bian Wu, Paul Hansma, Xifeng Yan, Kenneth S. Kosik, Kristofer Bouchard

## Abstract

Understanding how neuronal signals propagate in local network is an important step in understanding information processing. As a result, spike trains recorded with Multi-electrode Arrays (MEAs) have been widely used to study behaviors of neural connections. Studying the dynamics of neuronal networks requires the identification of both excitatory and inhibitory connections. The detection of excitatory relationships can robustly be inferred by characterizing the statistical relationships of neural spike trains. However, the identification of inhibitory relationships is more difficult: distinguishing endogenous low firing rates from active inhibition is not obvious. In this paper, we propose an *in silico* interventional procedure that makes predictions about the effect of stimulating or inhibiting single neurons on other neurons, and thereby gives the ability to accurately identify inhibitory causal relationships. To experimentally test these predictions, we have developed a Neural Circuit Probe (NCP) that delivers drugs transiently and reversibly on individually identified neurons to assess their contributions to the neural circuit behavior. With the help of NCP, three inhibitory connections identified by our *in silico* modeling were validated through real interventional experiments. Together, these methods provide a basis for mapping complete neural circuits.

## 1 Introduction

As proposed by D. O. Hebb [1] a “cell assembly” is a network of neurons that is repeatedly activated in a manner that strengthens excitatory synaptic connections. An assembly of this sort has a spatiotemporal structure inherent in the sequence of activations, and consequently strong internal synaptic strengths, which distinguish them from other groups of neurons. Although assemblies of this sort can be defined in numerous ways, one approach is to identify statistically significant time-varying relationships among simultaneously recorded neurons from the spike trains [2–6]. Obtaining these neural activity measurements requires recording from many neurons in parallel that can be spatially localized and temporally resolved at sub-millisecond time scales [7]. Widely used approaches for recording from multiple neurons such as calcium imaging and voltage sensitive dyes as a proxy for electrical activity or multiple implanted micro electrodes do not satisfy all these requirements. Novel instrumentation is required to meet the challenge of drawing complete neural circuits.

Dissociated neurons can self-organize, acquire spontaneous activity, and form networks according to molecular synaptogenic drivers that can be visualized and probed with multi-electrode arrays (MEAs). The work presented here utilizes MEAs to record signals sub-millisecond time resolution and precise spatial localization. The Neural Circuit Probe (NCP) uses mobile probes for local chemical delivery to a neural circuit of cultured neurons on a commercial MEA with 120 electrodes. Local drug delivery transiently and reversibly modulates the electrical behavior of individually identified neurons to assess their contributions to the circuit behavior. The dynamics of neuronal networks require both excitatory and inhibitory signals. Excitatory cells alone cannot generate “cell assemblies” because such interconnections would only lead to more excitation. A balance between excitatory and inhibitory neurons ensures the stability of global neuronal firing rates while allowing for sharp increases in local excitability which is necessary for sending messages and modifying network connections [8]. In a neuronal network described in terms of correlations among statistically significant time-varying relationships among the spike trains of simultaneously recorded neurons, the detection of excitatory relationships can be inferred based upon correlations between spikes with constant latencies that approximate synaptic transmission [9,10]. However, the identification of inhibitory relationships is more difficult: distinguishing endogenous low firing rates from active inhibition is not obvious.

In this paper, we demonstrate that tools from statistical inference can predict functionally inhibitory synaptic connections and show how inhibition propagates in a network to affect other neurons. We first fit a Generalized Linear Model (GLM) to spike trains recorded from neurons in hippocampal cultures, and inferred effective interactions between these neurons. We then used the fitted model to perform simulated *in silico* experiments in which we simulated the effect of silencing individual neurons in a network on the activity of other neurons. We tested the predictions from these simulated silencing experiments by performing real experiments in which we applied Tetrodotoxin (TTX) to silence neurons and thereby validated our computational approach toward the detection of inhibitory interactions

## 2 Methods

### 2.1 Cell culture

Commercial MEAs (Multi-electrode arrays) were sterilized with UV irradiation (for ¿ 30 minutes), incubated with a poly-D- or poly-L-lysine (0.1 mg/ml) solution for at least one hour, rinsed several times with sterile de-ionized water water and allowed to dry before cell plating. The culture chamber surrounding the MEA was 25 mm in diameter and filled with 1 ml of cell culture media. Cell cultures were prepared in two stages. This was done to allow glia to proliferate and become confluent in the area of the electrodes (1st plating) and for neurons to grow within a substrate of confluent glia (2nd plating). Unless otherwise stated, cells were cultured at 125,000 cells per dish. Mouse hippocampal neurons were used for all experiments described here. All mice were in a C57BL/6 genetic background and male mouse pups were used for all cell cultures. Mouse pups were decapitated at P0 or P1, the brains were removed from the skulls and hippocampi were dissected from the brain [11]. After one week, cultures were treated with 200 uM glutamate to kill any remaining neurons followed by a new batch of cells added at the same density as before. Cultures were grown in a tissue culture incubator (37°C, 5% CO2), in a medium made with Minimum Essential Media with 2 mM Glutamax (Life Technologies), 5% heat-inactivated fetal calf serum (Life Technologies), 1 ml/L of Mito+ Serum Extender (BD Bioscience) and supplemented with glucose to an added concentration of 21 mM. All animals were treated in accord with University of California and NIH policies on animal care and use.

### 2.2 Electrophysiology

Most recordings were done in cell culture medium so as to minimally disturb the neurons. In some cases we instead used an extracellular solution containing (in mM) 168 NaCl; 2.4 KCl; 10 HEPES; 10 D-glucose; 1.8 CaCl2; and 0.8 mM MgCl2. Pipette solution contained (in mM): 140 potassium gluconate; 4 CaCl2; 8 NaCl; 2 MgCl2; 10 EGTA; 2 Na2ATP and 0.2 Na2GTP. The pH was adjusted to 7.4 with KOH. The osmolality of external and internal solutions was adjusted to 320 mosmol. Salts were obtained from Sigma-Aldrich or Fluka; TTX was obtained from Ascent Scientific. Recordings were done using MultiChannel Systems MEA 2100 acquisition system. Data were sampled at 20 kHz and post-acquisition bandpass filtered between 200 and 4000 Hz. Recordings were done at 290 to 340 C. All recordings were done on neurons at 7-30 days in vitro (DIV). Data recordings were typically 3.5 to 5 minutes long. Recording duration was typically kept short to minimize the effects of removing MEAs from the incubator. Drug manipulations were done with a custom built instrument that allowed us to apply drug locally.

### 2.3 Spike sorting

For each MEA recording, we first removed redundancy propagation signals [12] and then did spike sorting [13]. Extracellular signals were band pass filtered using a digital 2nd order Butterworth filter with cutoff frequencies of 0.2 and 4 kHz. Spikes were then detected and sorted using a threshold of 6 times the standard deviation of the median noise level.

The data in Fig 3a were gathered in one recording session and each “unit” corresponds to one spike train after the spike sorting algorithms were applied on the raw data. However, the data in Fig 3d and Fig 3g were gathered in several recording sessions. So, the labels of units could be inconsistent in different recording sessions after the spike sorting algorithms were applied. Hence, to make the data consistent across different recording sessions, for these two datasets, we merged the spike trains from the same electrode as one unit.

### 2.4 A pipeline to identify and validate putative inhibitory connections

We used a novel pipeline to first identify putative inhibitory connections from spike trains and then validate them with a Neural Circuit Probe (NCP) that we built. Mouse hippocampal neurons were dissociated and plated on a multi-electrode array (MEA). To begin with, as shown in Fig 1a, their spontaneous spiking activity was modeled using a Generalized Linear Model (GLM) in which the outcome is a zero or one (spike or no spike) random variable and single neurons generate spikes according to a Poisson process. The rate of this process was determined by the spikes from other neurons. Parameters of the GLM were fit using a gradient descent algorithm to minimize the negative log likelihood of the recorded spike trains.

We next conducted *in silico* interventional experiments to identity inhibitory connections as shown in Fig 1b. Single neurons were silenced or activated in *silico* and then these data were used to infer predicted effects on connectivity using the fixed parameters from the GLM as determined above. The procedure for running the *in silico* interventional experiment was as follows. First, we selected one neuron as our interventional target. Throughout the simulation experiment, the state of this neuron was fixed to either 0 (silenced) or 1 (activated). Then, for all the other neurons, we ran the GLM with the inferred parameters to get the probabilities of seeing a spike at the next time point. Each probability represented how likely it was for a neuron to generate a spike at the next time point. Given the probability, we sampled a binary value (0 or 1) from a Bernoulli distribution as the state of the neuron for the next time point, where 0 refers to no spike and 1 means spike. We continued doing this to generate simulated recordings one time point at a time until a desired length *T* had been reached, where *T* is the number of time points in our *in silico* interventional recording. To find inhibitory connections, we investigated the generated simulated data to find those neurons that were negatively correlated (Pearson correlation coefficient) with the intervention taken on the target neuron. These neurons were considered as potentially inhibited by the interventional target.

Finally, we conducted real TTX delivery experiments to validate the putative inhibitory connections predicted from the *in silico* interventional experiments as shown in Fig 1c. In these experiments, TTX was delivered using the NCP as a delivery system. The NCP delivered TTX in a manner highly localized to a single electrode and in sufficiently low concentration that its potency dropped below threshold once it diffused beyond a single electrode. The NCP can detect increased impedance as the probe approached the cell and therefore allowed us to deliver TTX as close as possible to the cell without directly contacting the cell. Each TTX delivery resulted in the rapid onset of complete silencing of the neuron to which it was applied. As a result, putative inhibitory connections were validated when we observed activation of an inhibited neuron for a duration that approximated the time of TTX-induced silencing.

### 2.5 Generalized Linear Model

We used GLM to model the spiking of neurons. Let *m* denote the number of neurons being recorded and *x_i,t_* be the number of spikes of neuron *i* at time *t*. We assume *x_i,t_* is drawn from a Poisson distribution with rate λ**i,t** which is written as

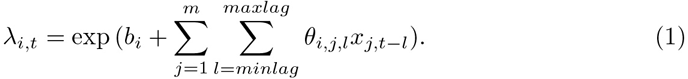

where *b_i_* is a parameter controlling the spontaneous firing rate of neurons *i* and *θ_i,j,t_* denotes the effective interaction from neuron *j* to neuron *i* at time lag *l*. We assume that the firing rate of neurons *i* depends on the activities of all neurons in a history window that spans from time *t – maxlag* to time *t – minlag*, where *minlag* and *maxlag* are the minimum and maximum time lags we consider.

Given Eq. 1 for the firing rate of individual neurons, the likelihood for the observation of neuron *i* at time *t*, *L_i,t_* is

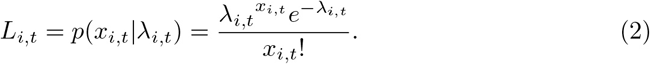

In spike train data with one millisecond time bin, there are at most one spike at any time point and therefore *x_i,t_* takes the value of 0 or 1. Hence, the log-likelihood is

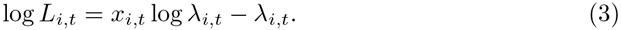

The log-likelihood for all the observations in a recording with length *T* is

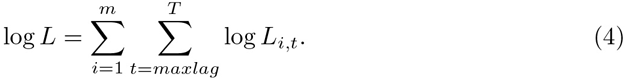

**Fig 1.**
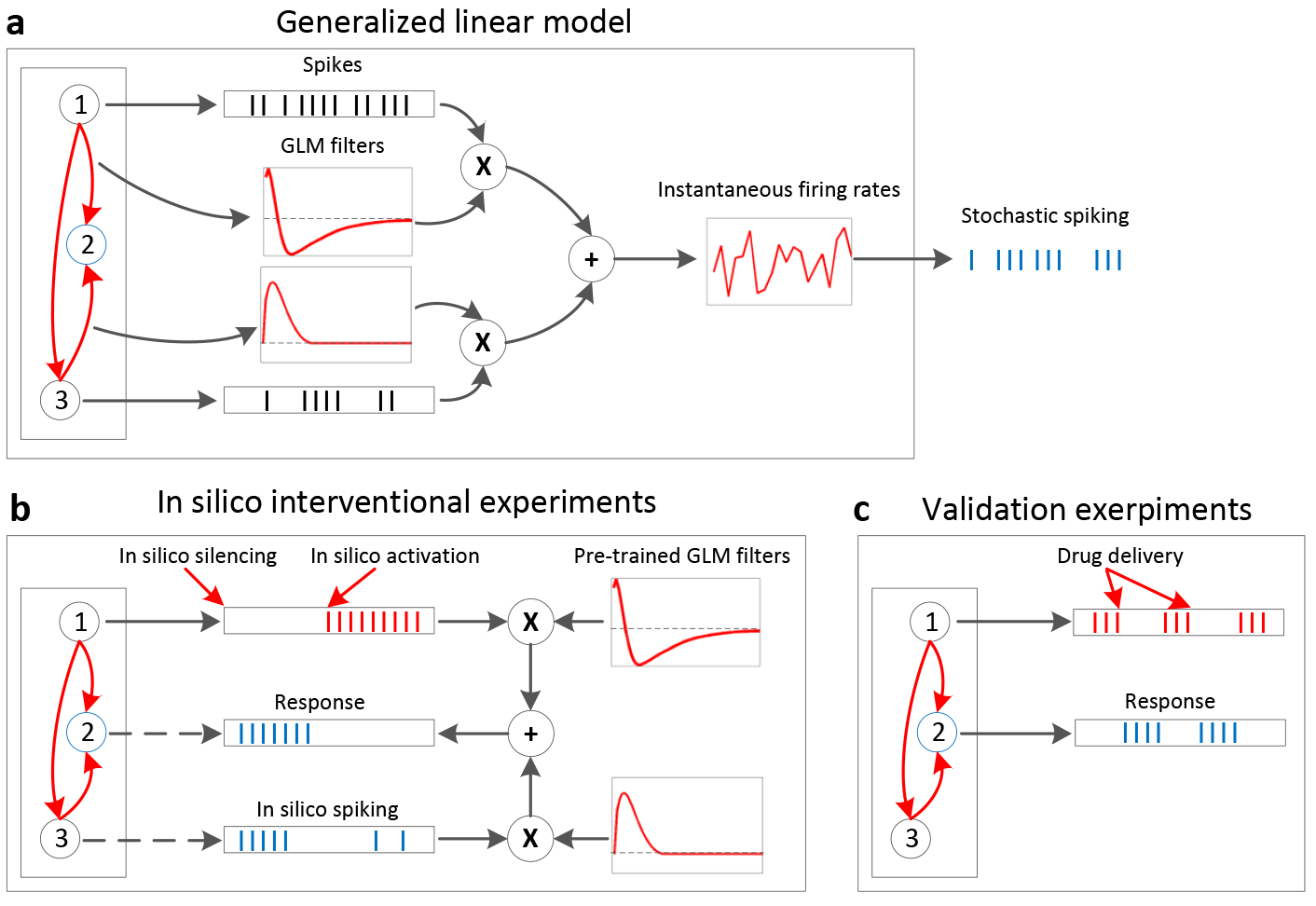
An overview of the procedure that we used to identify and validate direct and indirect inhibitory connections. (a) A Generalized Linear Model (GLM), in which the firing of a neuron is modeled as determined by the spikes from other neurons, was used. Filters of the GLM were inferred from a training recording of spontaneous firings. (b) In silico experiments were conducted by performing simulated interventions on a neuron and generating simulated responses with pre-trained GLM filters. Putative inhibitory connections were then identified by comparing the simulated interventions and responses. (c) Real drug delivery experiments were conducted to validate the putative inhibitory connections.

The model described above includes too many parameters and there is nothing in the model that ensures the inferred parameters to vary smoothly with time, something that isas expected from interactions between pairs of neurons. Furthermore, the model has too many parameters and this might cause problems for robustly inferring them. To ensure the smoothness of the filters, instead of directly using a history window of spikes in the model, following [14], we use their filtered versions that are created by convolving with several cosine bumps. To minimize the number of fitting parameters and prevent overfitting, we add an *L* — 1 regularizer to the likelihood. These remedies are described further below.

We first design *p* cosine basis functions where the *l^th^* cosine basis function can be written as:

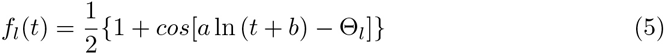

for all times *t* such that satisfy

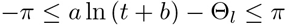

and

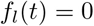

outside the interval defined above. The values of *a, b* and Θ*_l_* are manually chosen. One of the factors to be considered when choosing their values is the locations where the peaks of the bumps occur. During experiments, we used pairwise cross-correlations to determine the locations of the peaks.

In the *naive* GLM without the basis functions, for neuron *j*, we used a history window of spikes to model its influence on other neurons. Now the raw spikes are convolved with *p* cosine basis functions to get the filtered versions, of which the *l^th^* value is calculated as follows:

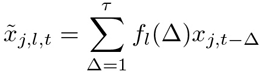

where *τ* is the length of the history window that is covered by the cosine basis functions. Eq. (1) is rewritten as:

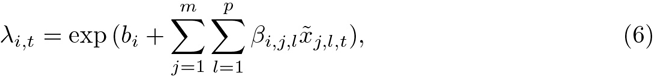

where *β_i,j,l_* is the weight of the *l^th^* basis function for the influence from the neuron *j* to neuron *i*.

As mentioned above, to prevent overfitting, we added an *L*1 regularization term to penalize non-zero filter parameters. The loss function we want to minimize is rewritten as

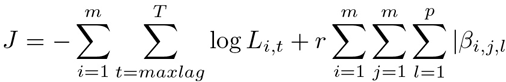

where *r* is the regularization constant. The value of *r* is decided by doing 10-fold cross validation on a spontaneous firing recording of 60 seconds. We used the Area Under the Receiver Operating Characteristic curve (AUC-ROC) as our metric to evaluate the performance of the fitted model to do predictions on future spikes given previous spiking histories.

### 2.6 *in silico* interventional experiments

To identify inhibitory connections from an ensemble of neurons, one straightforward way is to investigate the GLM filters obtained by fitting the spike trains, as these filters represent the relations of neurons captured by GLM. However, the inhibitory effects among neurons can be rather complex than obvious, and simply using the GLM is usually not enoughsufficient. For example, two of the inhibitory connections we identified in this study were not observable from the their corresponding GLM filters, but became obvious once interventions were applied. Therefore, in this study, we have proposed a method to conduct in *silico* interventional experiments which could discover hidden inhibitory connections by running simulated experiments.

To cold start the simulated experiment, we used a history window of length *τ* with none spiking states (i.e., all the neurons take the value 0 in a time window of *τ*). The instantaneous firing rate of neuron *i* at time *t* was calculated according to E.q. (6) in Methods section. Therefore, the probabilities of seeing and not seeing a spike are

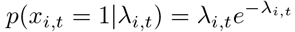

and

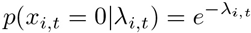

Because in our setting, there is at most one spike in the one millisecond time bin, *x_i,t_* can only take the value of 0 or 1. However, if we run simulated experiments by directly sampling from a Poisson distribution, the value *x_i,t_* takes could be arbitrary instead of binary. Hence, we normalize the probability of getting a spike at time point *t* as

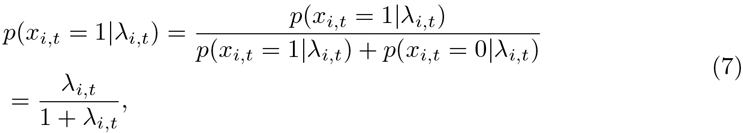

For neuron *i* at time point *t*, we generate the simulated sate by sampling a value from a Bernoulli distribution with the probability of Eq. (7).

During the *in silico* interventional experiments, we selected one neuron as our interventional target and fixed its state to be either 0 (silenced) or 1 (activated). Then, the responses from other neurons were gathered and compared with the intervention taken on the target neuron by calculating their Pearson correlation coefficients. Those neurons that were negatively correlated with the intervention were considered as potentially inhibited by the interventional target. The algorithm is shown in Algorithm 1.

#### Algorithm 1 Identifying Putative Inhibitory Connections

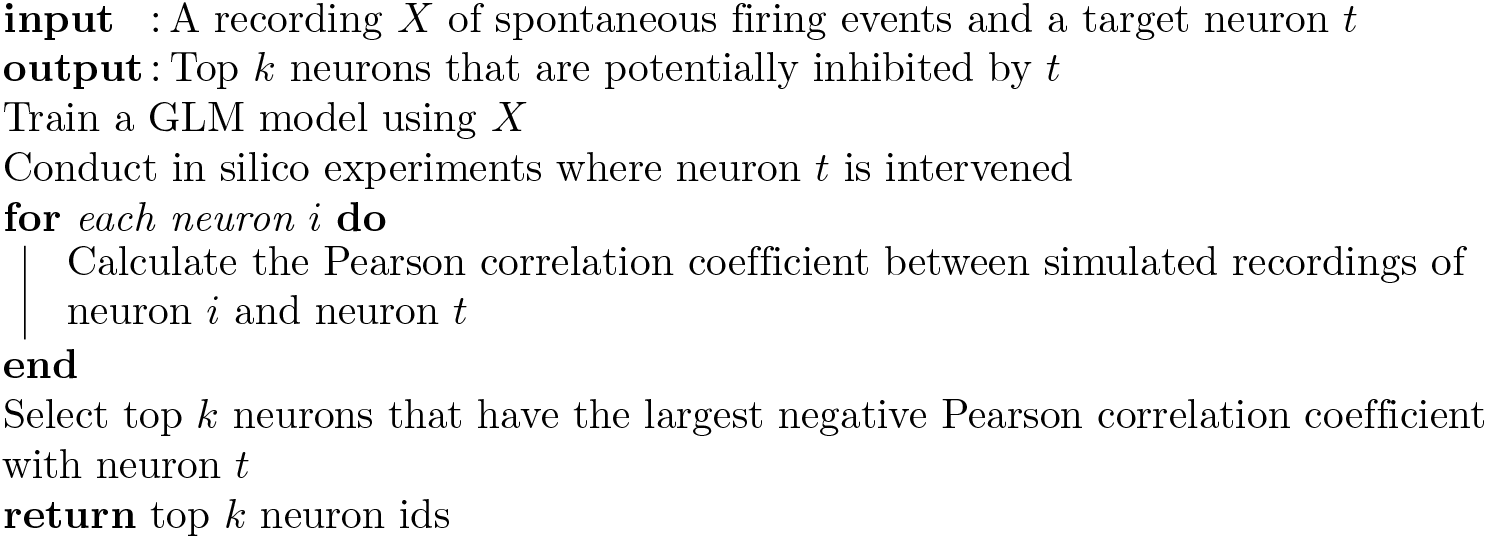

### 2.7 Instrumentation for validating putative inhibitory connections

Identification of single cell contributions to a neuronal circuit requires precise access to and control over functionally identified cells. To accomplish this goal we built a neural circuit probe (NCP) consisting of (1) a head unit that accepts various probes, (2) an integrated perfusion chamber plus light ring illumination system, (3) a probe control system with computer interface which implements a simple feedback system for an automated approach, and (4) a commercial MEA (MEA2100, Multi Channel Systems) mounted to a custom X-Y translation stage (Fig 2).

The NCP controller uses proportional and integral feedback control to position the various probes, and can accept a variety of input signals, such as ion current used here. An amplifier is located on the head unit that amplifies the current signal before going to the controller. The NCP software allows the operator to engage and disengage the probe using feedback. It is also used to control the location of the MEA stage beneath the probe, allowing the operator to position the probe above neurons of interest. A pneumatic control system attached to the probe regulates a pressure line for chemical delivery (Fig 2a, Fig 2b). An integrated pressure sensor, connected to the MEA data acquisition system, measures the duration and magnitude of pressure for temporal alignment with the MEA signal.

Local targeted drug delivery with the NCP can be used to modify their electrical behavior. This was done with small pipettes typically with inner diameters of 1-2 microns. In this example (Fig 2c, Fig 2d) we applied the Na+ channel blocker tetrodotoxin (TTX, 500 nM) to induce a temporary and reversible cessation of activity from that cell. Thus with the NCP we can do targeted drug delivery with high spatial resolution.

**Fig 2.**
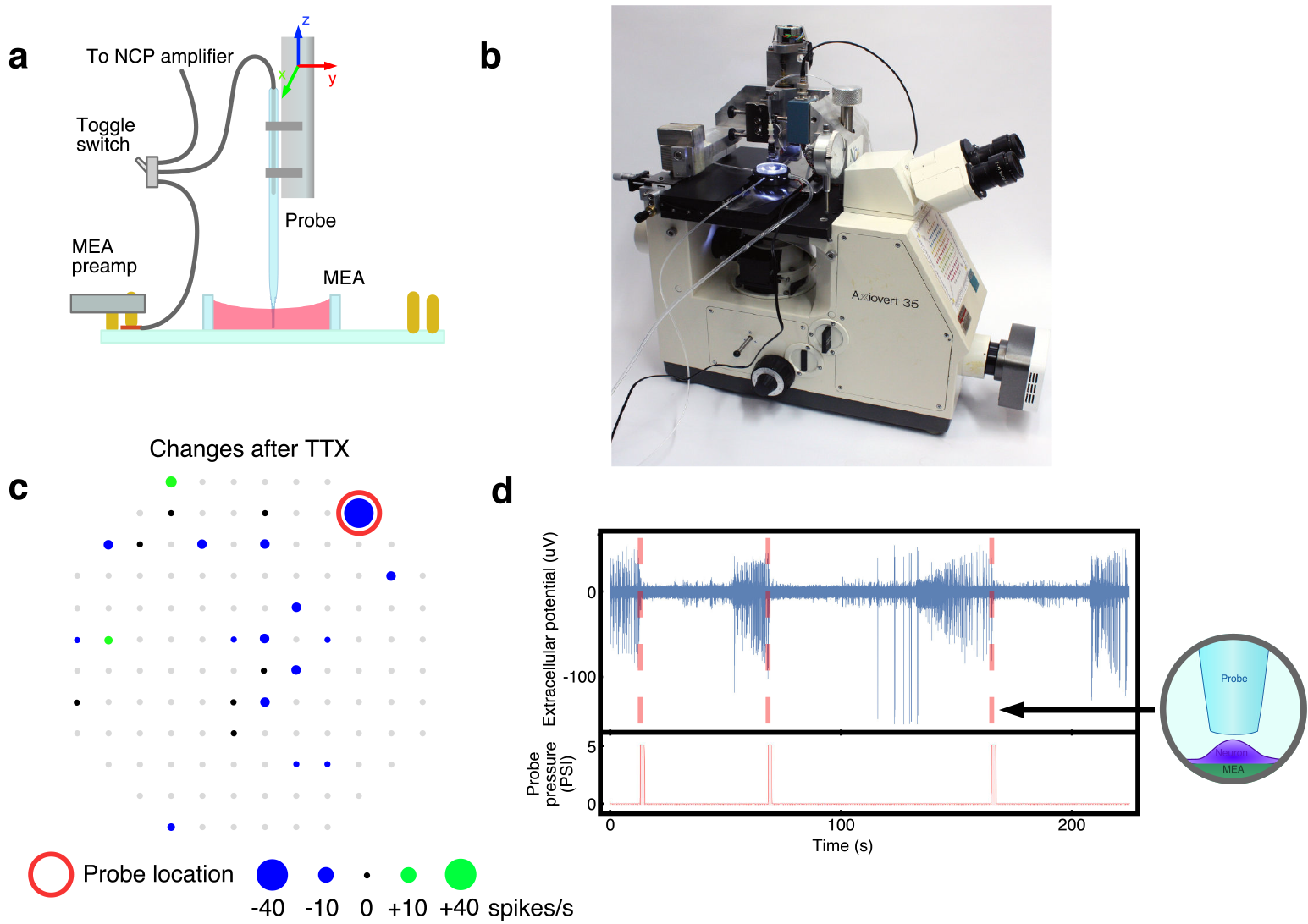
Illustration of the Neural Circuit Probe (NCP) and real drug delivery experiments to validate putative inhibitory connections. (a) Schematic drawing of the key components. The probe is positioned in x and y to center it in the field of view of the microscope. Then the MEA is translated in x and y to bring a target neuron directly under the probe. Finally the probe is automatically lowered, with ion conductance feedback, to just above, but not touching, the neuron.(b) Overview of the NCP situated on an inverted microscope. (c) The changes of firing rates at all electrodes before and after TTX application. Gray dots are electrodes with no spiking activities recorded. Black dots are electrodes with no spiking rate changes. When we blocked spiking at the specific electrode (red circle) it had widespread secondary effects on the firing rates observed at other MEA electrodes. Though the firing rate decreased for many electrodes (blue dots), for two electrodes it increased (green dots). (d) A transient increase of probe pressure delivered TTX (500 nM), which reversib y blocked spiking activity, with high spatial resolution. This process was repeated 3 times.

## 3 Results

### 3.1 Identifying Putative Inhibitory Connections

Following the aforementioned procedure, a recording with spontaneous activity from 17 units over a duration of 20 seconds divided into one millisecond time bins was used to fit the GLM model (Fig 3a). Each unit corresponded to a spike train after spike sorting and removal of the redundancy inherent in propagation signals [12]. Then unit 10 was chosen as the in silico interventional target, i.e. it was fixed in a silent state (no spikes at all times) for 10 seconds and then fixed for 10 seconds in an active state (continuous spiking). Simulations with the fitted GLM identified five units with the highest probability to be inhibited by unit 10 (Fig 3c). The strongest candidate for inhibition by unit 10 was unit 8. Note that the filters from the fitted GLM also suggested that the connection from unit 10 to unit 8 was predominantly inhibitory (Fig 3b).

**Fig 3.**
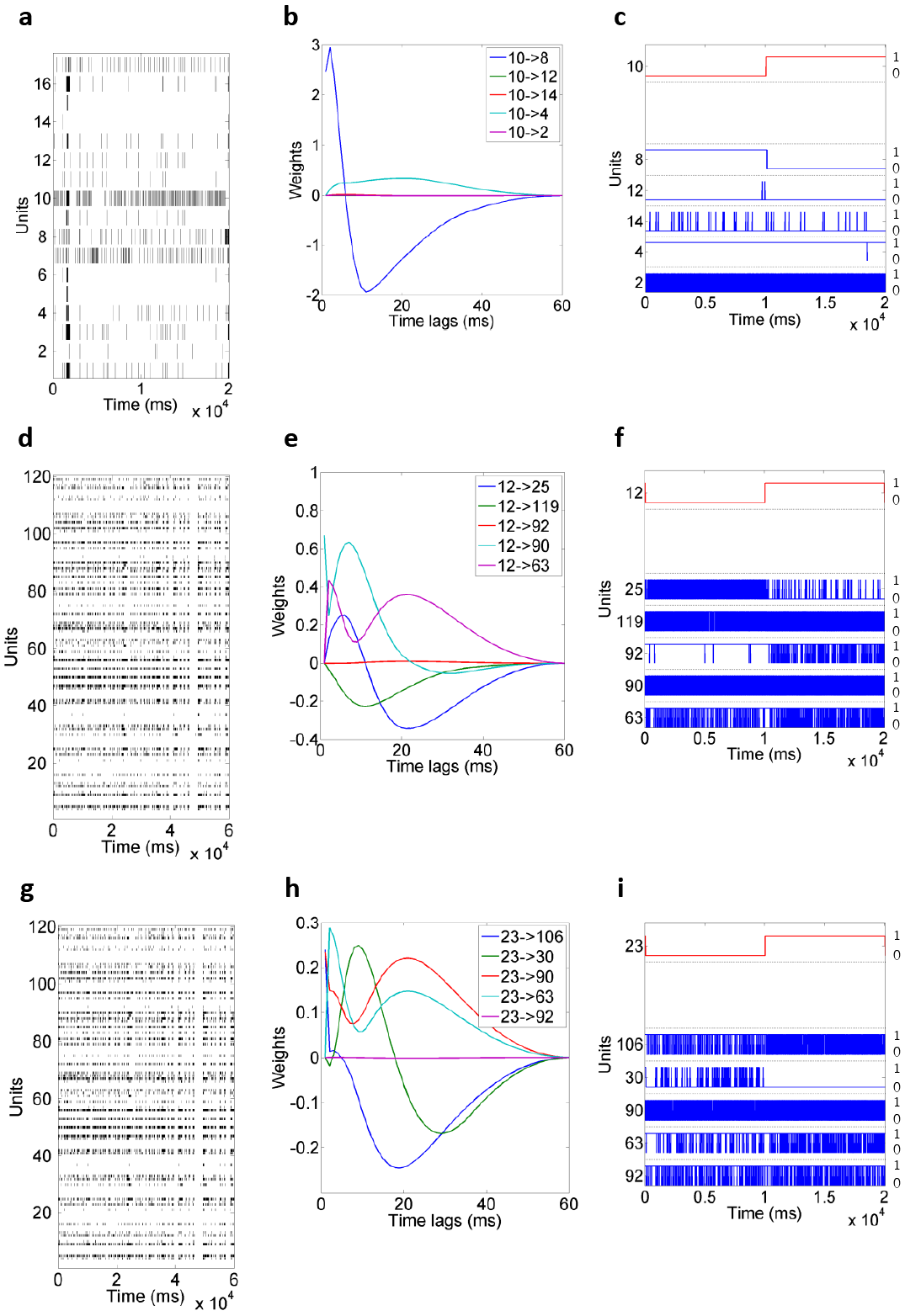
Real data examples of the procedure that we used to identify putative inhibitory connections. (a) A training recording of 17 units for a duration of 20 seconds which were divided into 1 millisecond time bins. The black bars represent spikes. (b) Filters of the GLM inferred from the training recording. Note that at different time lags, the strength of the connection between two units is also different. (c) Simulated data for top 5 units that were negatively correlated with the intervened unit 10. The red and blue lines represent the instantaneous firing rates for the simulated recordings. The labels on the left of the y-axis represent the unit numbers and the labels on the right represent the range of the instantaneous firing rates (0 to 1). Note that when unit 10 was changed from silent state to active state, conversely, unit 8 changed to silent state from active state, which implied a putative inhibitory connection. (d) A training recording of 120 electrodes for a duration of 60 seconds which were divided into 1 millisecond time bins. (e) Filters of the GLM inferred from the training recording. (f) Simulated data for top 5 units that were negatively correlated with the intervened unit 12. (g) A training recording of 120 electrodes for a duration of 60 seconds which were divided into 1 millisecond time bins. (h) Filters of the GLM inferred from the training recording. (i) Simulated data for top 5 units that were negatively correlated with the intervened unit 23.

Additional *in silico* experiments on another cell culture were also conducted to identify putative inhibitory connections by following the aforementioned procedure. For these experiments, we used a 60 second recording of spontaneous firing events (Fig 3d) to fit a GLM. The GLM parameters for the connections from unit 12 to five units are shown in Fig 3e. We calculated the Pearson correlation coefficients between the in silico intervention on unit 12 and simulated responses of every other neuron. The top five negatively correlated units were chosen and investigated (Fig 3f). In another example, we chose unit 23 as the in silico interventional target. Similarly, Fig 3h shows the GLM parameters for the connections from unit 23 to five other units and Fig 3i shows the top five units that were identified as candidates for inhibition by unit 23.

### 3.2 Validation of Inhibitory Connections

Given putative inhibitory connections identified in the first example (Fig 3c), to validate experimentally that unit 8 was an inhibitory target of unit 10, TTX was delivered four times on unit 10 (Fig 4a) using the neural circuit probe as a delivery tool. The instrument delivered TTX in a manner highly localized to a single electrode and in sufficiently low concentration that its potency dropped below threshold once it diffused beyond a single electrode. Each TTX delivery resulted in the rapid onset of complete silencing of the neuron to which it was applied. Delivery of TTX to the electrode corresponding to unit 10 resulted in the activation of unit 8 and activation of the target neuron for a duration that approximated the time of TTX-induced silencing. These experimental data clearly demonstrated that the top inhibitory connection (from 10 to 8) predicted by our simulated experiment was validated by the actual TTX delivery experiment.

**Fig 4.**
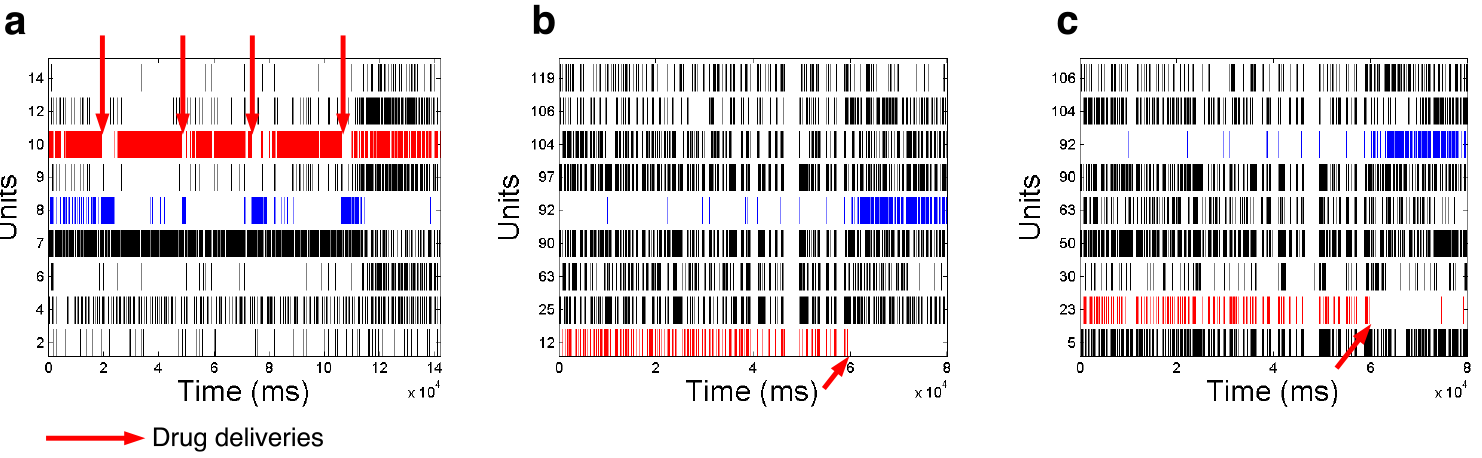
Real TTX experiments to validate putative inhibitory connections. (a) Real TTX experimental recording where unit 10 was silenced 4 times by delivering TTX. Unit 8 rebounded every time unit 10 was silenced, which indicated an inhibitory connection from 10 to 8. (b) Real TTX experimental recording where unit 12 was intervened. (c) Real TTX experimental recording where unit 23 was intervened.

In the second example, 92 was a strong candidate for inhibition by unit 12. To validate this inferred connection experimentally, we delivered TTX to unit 12 and, as predicted, observed an inhibitory effect from unit 12 to unit 92 (Fig 4b). It’s also worth mentioning that even though unit 92 is not the top 1 candidate predicted by our *in silico* interventional experiments, it’s within the top 5 predictions out of 120 possible units. This shows that the in silico interventional experiments could give accurate predictions of putative inhibitory connections. In the final example, we also delivered TTX to unit 23 and observed rebound of firing on unit 92 (Fig 4c) which was predicted by the in *silico* interventional experiments (Fig 3i).

### 3.3 Indirect connections

The inhibitory connections identified in this study may not be direct. A unit could be causing inhibitory effects on another unit through a third unit. To study the possibilities of inhibitory connections, we have revisited the three examples of inhibitory connections validated in Fig 4. For each example, we introduced a third unit and convolved the GLM filters of the two connections in a potential inhibitory connection. Fig 5 shows the convolution outputs that exhibited inhibitory effects. To understand the inhibitory effects from unit 10 to unit 8, we show three possible cases where unit 10 could cause an inhibitory effect on unit 8 through a third unit (Fig 6a).

**Fig 5.**
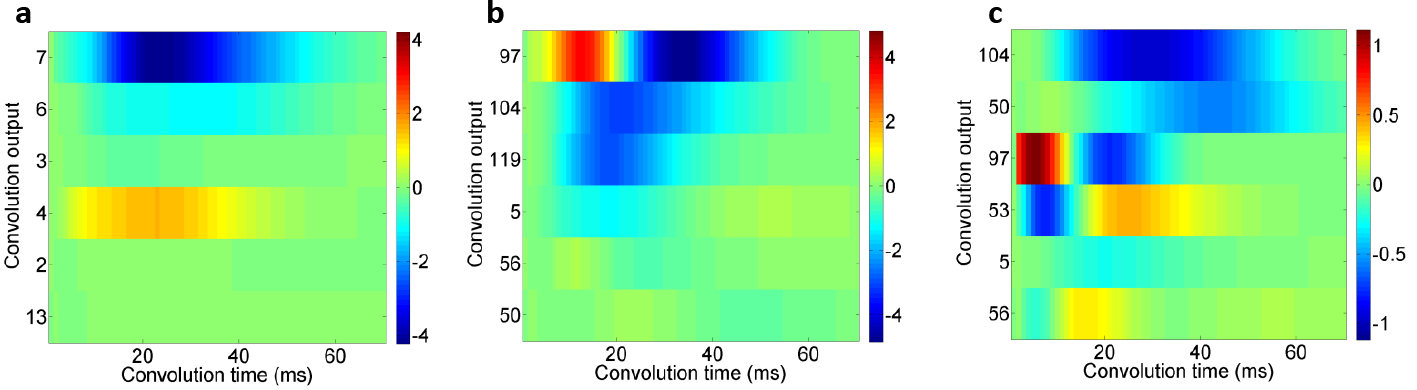
Convolutions of the GLM filters from indirect connections. (a) Convolution of the GLM filters from the connection 10 → *m* and *m* → 8, where *m* (*y*-axis) is an intermediate unit. The convolutions when *m* is 10 or 8, which indicates a direct connection, are omitted. (b) Convolution of the GLM filters from the connection 12 → *m* and *m* → 92, where *m* (*y*-axis) is an intermediate unit. (c) Convolution of the GLM filters from the connection 23 → *m* and *m* → 92, where *m* (*y*-axis) is an intermediate unit.

**Fig 6.**
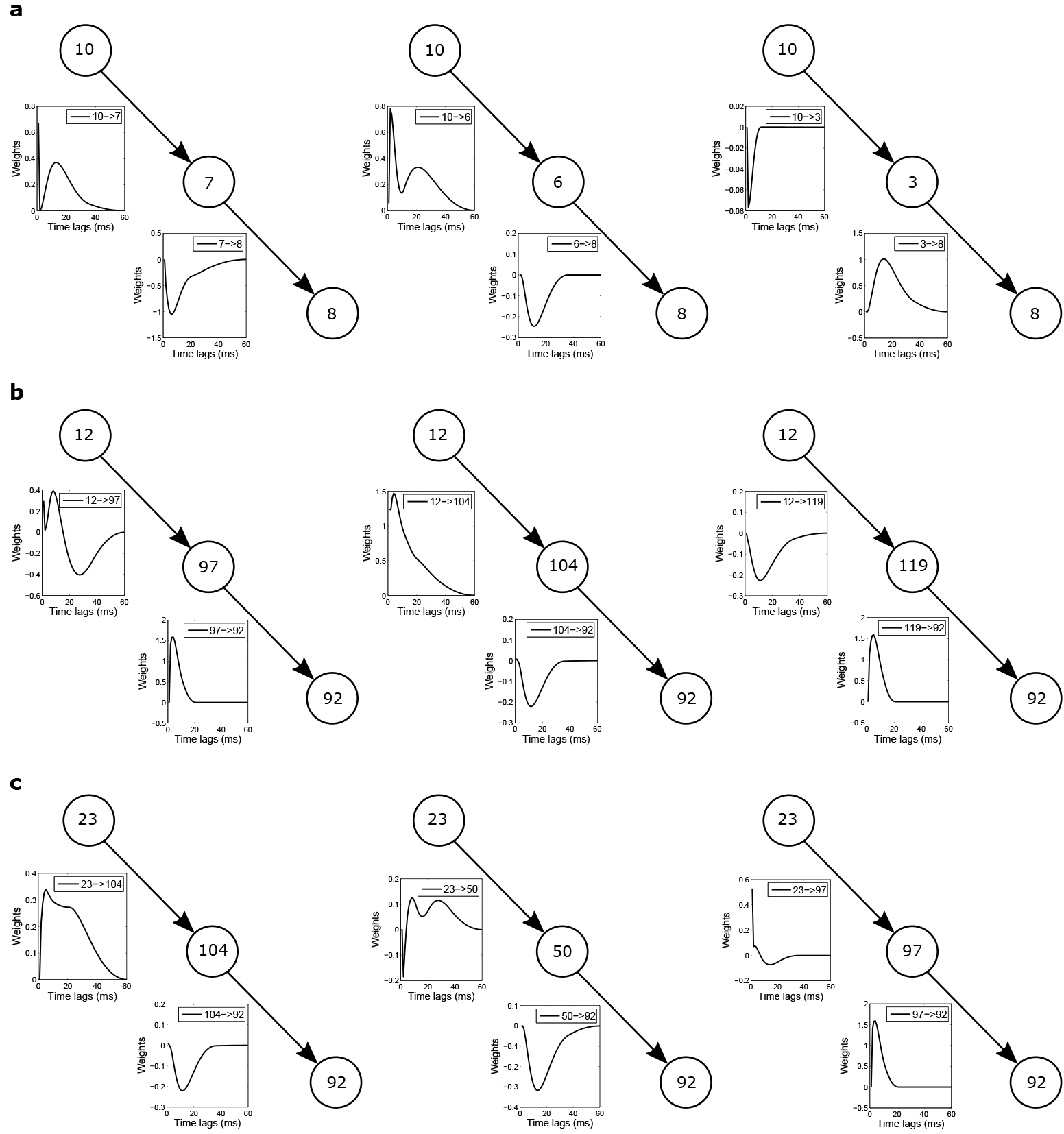
Indirect inhibitory connections. (a) Three possible cases where unit 10 has an inhibitory influence on unit 8through a third unit. The first case consists of an excitatory connection (10 → 7) and an inhibitory connection (7 → 8).The second case consists of an excitatory connection (10 → 6) and an inhibitory connection (6 → 8). The third case consists of an inhibitory connection (10 → 3) and an excitatory connection (3 → 8). (b) Three possible cases where unit 12 has an inhibitory influence on unit 92 through a third unit. The first case consists of a predominantly inhibitory connection (12 → 97) and an excitatory connection (97 → 92). The second case consists of an excitatory connection (12 → 104) and an inhibitory connection (104 → 92). The third case consists of an inhibitory connection (12 → 119) and an excitatory connection (119 → 92). (c) Three possible cases where unit 23 has an inhibitory influence on unit 92 through a third unit. The first case consists of an excitatory connection (23 → 104) and an inhibitory connection (104 → 92). The second case consists of an excitatory connection (23 → 50) and an inhibitory connection (50 → 92). The third case consists of a predominantly inhibitory connection (23 → 97) and an excitatory connection (97 → 92).

The second example shown in Fig 3 and Fig 4 illustrates an important feature that the in silico experiments offers in describing how signals propagate in the network. In this example the inhibitory effects from 12 to 92 is not obviously manifested in the filters shown in Fig 3e, i.e. the magnitude of the curve representing the connection from 12 to 92 is not as significant as others. However, this inhibitory effect is ranked high according to the negative Pearson correlation score given the simulated experimental results. One explanation for this is the indirect connections among units. It may be the case that unit 12 is not directly inhibiting unit 92, but it could cause an inhibitory effect through other units.

To explore this possibility further, we show three possible indirect inhibitory connections from unit 12 to unit 92 (Fig 6b). Each indirect connection consists of a predominantly excitatory connection and a predominantly inhibitory connection, which could cause a net inhibitory effect. Therefore, it supports the idea that the inhibitory effect from unit 12 to unit 92 were caused by indirect inhibitory connections.

As a final example, Fig 4c shows another inhibitory effect between pairs of neurons, in this case unit 23 to unit 92, as discovered from the *in silico* experiments on the fitted GLM and then validated by experiments. Similarly, we show three indirect inhibitory connections from 23 to 92 (Fig 6c).

## 4 Discussion

Understanding how neuronal signals propagate in local network is a prerequisite to understanding information processing in those networks. The ‘gold standard’ way to predict how the activity of one neuron influences another is through intracellular paired recordings along with pharmacologic probes. Using such intracellular recordings, one can establish the presence or absence of direct or indirect connections between pairs of neurons and thus to some degree predict how activity in one neuron affects the others. Inspired by the successes of this technology, we show here how it can be extended to larger networks of neurons using advanced mechatronic positioning of a probe over an array of electrodes with the Neural Circuit Probe. As a demonstration of the potential power of this device, we demonstrated its utility in testing the predictions of *in silico* modeling.

We first fitted a GLM model to spikes recorded from a culture using MEAs, then performed *in silico* experiments in which we silenced one of the units, and identified what other units will change their activity upon this inactivation. We then went back to the culture and silenced the same unit using TTX and observed that the inhibitory effects predicted by the *in silico* experiments showed up when TTX was applied.

The results presented here thus opened the door to using statistical models not only to characterize the statistics of neural spike trains or functional connectivity between neurons, but to make predictions about the response of the network to changes. Although using GLM to study the circuitry of a neuronal network is never going to be as accurate as intracellular recordings, the simplicity of fitting the model to data, and performing in silico experiments with this model, are great advantages that support the idea of using this approach to make educated guesses about the likely outcomes of manipulations to the network, i.e. offering a virtual culture, similar to a previous attempt to use GLMs to build a virtual retina. [15].

In using the GLM in neural data analysis, one typically assumes that a single neuron generates spikes via e.g. a Poisson process. The rate of this process is determined by the spikes from other neurons filtered by interactions that are inferred from data using convex optimization. The inferred model is then used for a variety of purposes that include evaluating the role of correlations in shaping population activity, for example, in the retina [14], the motor cortex [16,17], the functional connectivity between grid cells [18], or the relative influence of task related covariates on shaping neural responses in the parietal cortex [19]. Despite the widespread use of the GLM in neural data analyses, a potentially very powerful aspect of this class of models has been left unexplored: the ability of the GLM to make predictions about how a neuronal network responds to interventions. At the microcircuit level, this amounts to identifying meaningful interactions between pairs of neurons and using them to make predictions about how external manipulations of one or more neurons can affect the others. The main reason for the fact that GLMs have not been used for this purposes so far is that, in general, the ground truth about connectivity is not known and, therefore, it is not possible to compare the interactions inferred by GLM with the real ones. The results presented in this paper add a new dimension to how these statistical models can be used in neuroscience by showing that, although the relationship between individual synaptic interactions and those inferred by the GLM may not be known, the inferred connections can still be employed to make specific predictions about the functional connectivity of a neuronal network. Our results thus demonstrate how statistical models can be used to infer neuronal microcircuitry at a detailed level without using more complicated experimental techniques such as multi-unit intracellular recordings.

## Supplementary materials

**S1 Appendix. Choosing the regularization constant** The performance of the GLM was evaluated using a Receiver Operating Characteristic (ROC) curve, which plots true and false positive rates on different axes. The extent to which true positive rates exceed false positive rates is given by the area under the curve (AUC) and was our performance metric to evaluate how well the fitted model could predict future spikes given previous spiking histories. As we can see from Fig 7, the AUC-ROC stops increasing as we increase the regularization constant r to a certain point. We chose *r* (2.5 in our case) that gave us the best AUC-ROC.

**Fig 7.**
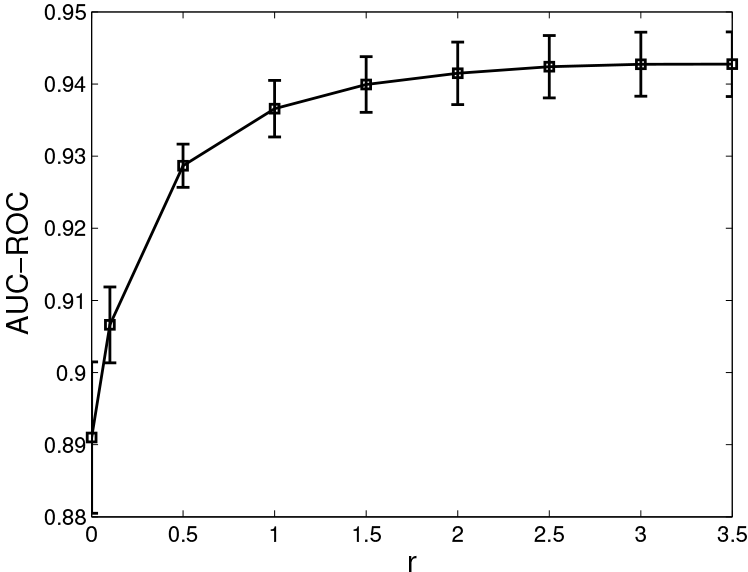
The AUC-ROCs with respect to *r*. For each value the regularization term has taken, we do a 10-fold cross validation and report the mean and variance.

**S2 Appendix. Additional functionality of the NCP** The NCP also incorporates a mobile electrode to measure extracellular potentials anywhere on the array, providing a higher spatial resolution of the signal versus the standard MEA (Fig 8). The mobile electrode consists of a 20 micron inner diameter micropipette with a 75 micron platinum electrode inserted in the top [20]. A single channel from the commercial MEA preamp is repurposed to measure and record from the mobile electrode. To use the commercial system, the repurposed channel was tapped into using a small piece of conductive film with an insulating layer on the backside, bypassing the electrode on the MEA, and inserting the mobile signal. This technique was necessary to validate the specific neuron that gave rise to the MEA signal of interest.

**Fig. 8.**
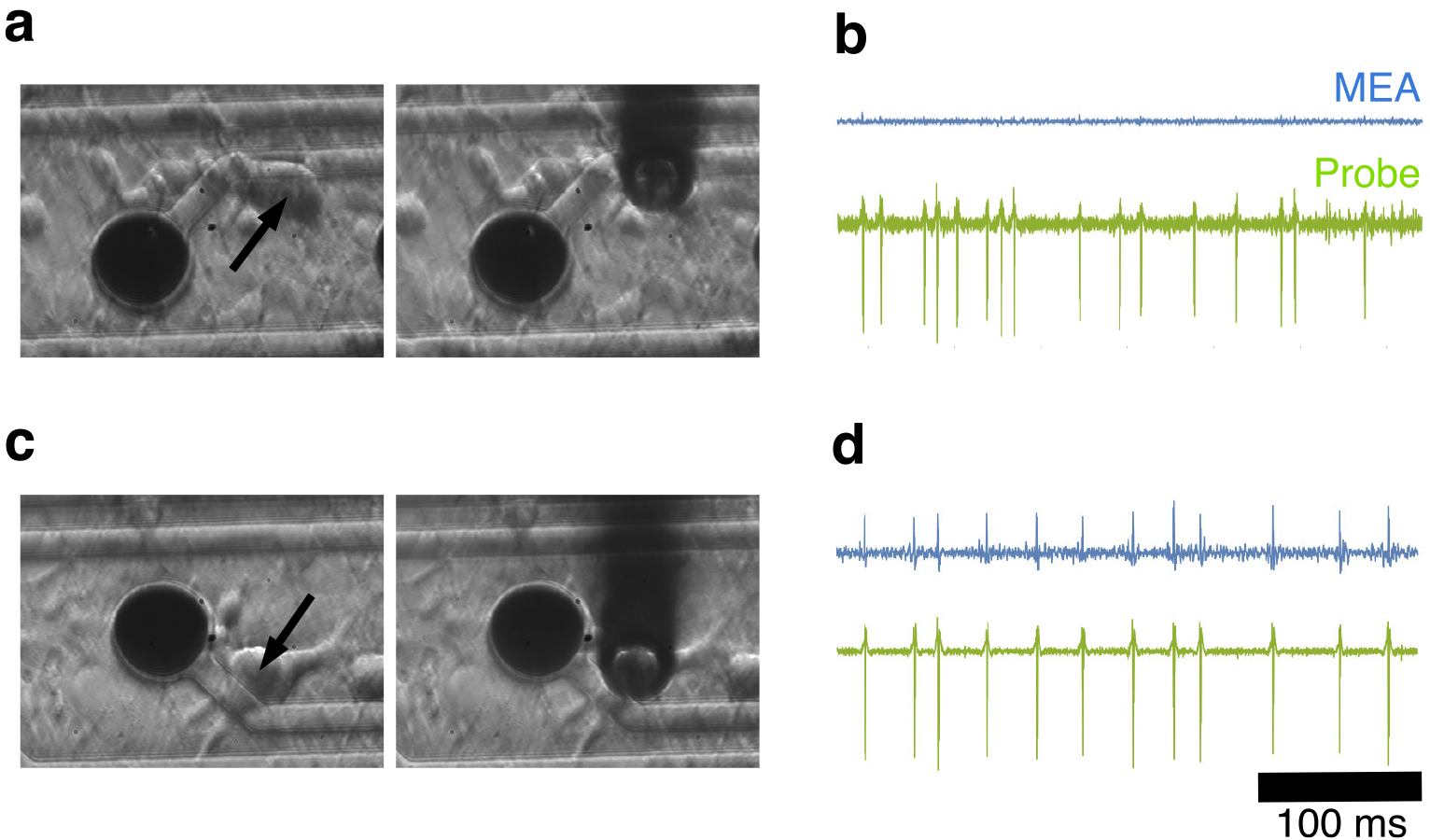
Monitoring electrical activity of a selected neuron. (a, b) No spiking activity was detected on the MEA for the neuron (arrow) closest to the MEA electrode (black circle) whereas a simultaneous recording from the mobile electrode shows robust spiking behavior (compare labelled traces in b). (c, d) The MEA electrode (black circle) closest to the neuron (c, arrow) showed spiking (d, top trace) but when the mobile electrode probe was positioned directly over the cell, the corresponding recording had a higher signal-to-noise ratio.

## Acknowledegment

This research was sponsored by the U.S. Army Research Laboratory and Defense Advanced Research Projects Agency under Cooperative Agreement Number W911NF-15-2-0056. The views, opinions, and/or findings contained in this material are those of the authors and should not be interpreted as representing the official views or policies of the Department of Defense or the U.S. Government. Additional support was also provided by the California NanoSystems Institute (CNSI). The funders had no role in the study design, data collection and analysis, decision to publish, or preparation of the manuscript.

## References

1. Hebb DO. The organization of behavior: A neuropsychological approach. John Wiley&Sons: 1949.

2. Keat J, Reinagel P, Reid RC, Meister M. Predicting every spike: a model for the responses of visual neurons. Neuron. 2001;30(3):803–817.

3. Pillow JW, Paninski L, Uzzell VJ, Simoncelli EP, Chichilnisky E. Prediction and decoding of retinal ganglion cell responses with a probabilistic spiking model. Journal of Neuroscience. 2005;25(47):11003–11013.

4. Stevenson IH, Rebesco JM, Miller LE, Körding KP. Inferring functional connections between neurons. Current opinion in neurobiology. 2008;18(6):582–588.

5. Stevenson IH, Rebesco JM, Hatsopoulos NG, Haga Z, Miller LE, Kording KP. Bayesian inference of functional connectivity and network structure from spikes. IEEE Transactions on Neural Systems and Rehabilitation Engineering. 2009;17(3):203–213.

6. Pillow JW, Ahmadian Y, Paninski L. Model-based decoding, information estimation, and change-point detection techniques for multineuron spike trains. Neural computation. 2011;23(1):1–45.

7. Gerstein GL, Kirkland KL. Neural assemblies: technical issues, analysis, and modeling. Neural Networks. 2001;14(6):589–598.

8. Jonas P, Buzsaki G. Neural inhibition. Scholarpedia. 2007;2(9):3286.

9. Cohen MR, Kohn A. Measuring and interpreting neuronal correlations. Nature neuroscience. 2011;14(7):811–819.

10. Ostojic S, Brunel N, Hakim V. How connectivity, background activity, and synaptic properties shape the cross-correlation between spike trains. Journal of Neuroscience. 2009;29(33):10234–10253.

11. Tovar KR, Westbrook GL. Amino-terminal ligands prolong NMDA receptor-mediated EPSCs. Journal of Neuroscience. 2012;32(23):8065–8073.

12. Tovar KR, Bridges DC, Wu B, Randall C, Audouard M, Jang J, et al. Recording action potential propagation in single axons using multi-electrode arrays. bioRxiv. 2017; p. 126425.

13. Quiroga RQ, Nadasdy Z, Ben-Shaul Y. Unsupervised spike detection and sorting with wavelets and superparamagnetic clustering. Neural computation. 2004;16(8):1661–1687.

14. Pillow JW, Shlens J, Paninski L, Sher A, Litke AM, Chichilnisky E, et al. Spatio-temporal correlations and visual signalling in a complete neuronal population. Nature. 2008;454(7207):995.

15. Bomash I, Roudi Y, Nirenberg S. A virtual retina for studying population coding. PloS one. 2013;8(1):e53363.

16. Truccolo W, Eden UT, Fellows MR, Donoghue JP, Brown EN. A point process framework for relating neural spiking activity to spiking history, neural ensemble, and extrinsic covariate effects. Journal of neurophysiology. 2005;93(2):1074–1089.

17. Rebesco JM, Stevenson IH, Körding KP, Solla SA, Miller LE. Rewiring neural interactions by micro-stimulation. Frontiers in systems neuroscience. 2010;4.

18. Dunn B, Mørreaunet M, Roudi Y. Correlations and functional connections in a population of grid cells. PLoS computational biology. 2015;11(2):e1004052.

19. Park IM, Meister ML, Huk AC, Pillow JW. Encoding and decoding in parietal cortex during sensorimotor decision-making. Nature neuroscience. 2014;17(10):1395–1403.

20. Claverol-Tinture E, Pine J. Extracellular potentials in low-density dissociated neuronal cultures. Journal of neuroscience methods. 2002;117(1):13–21.

